# Three-dimensional analysis of β-cell proliferation by a novel mouse model

**DOI:** 10.1101/659904

**Authors:** Shinsuke Tokumoto, Daisuke Yabe, Hisato Tatsuoka, Ryota Usui, Muhammad Fauzi, Ainur Botagarova, Hisanori Goto, Pedro Luis Herrera, Masahito Ogura, Nobuya Inagaki

**Author notes:** Corresponding author (Nobuya Inagaki).

## Abstract

Inducing β-cell proliferation could inhibit diabetes progression. Many factors have been suggested as potential β-cell mitogens, but their impact on β-cell replication has not been confirmed due to the lack of a standardized β-cell proliferation assay. In this study, we developed a novel method that specifically labels replicating β cells and yields more reproducible results than current immunohistochemical assays. We established a mouse line expressing the fluorescent ubiquitination-based cell cycle indicator (Fucci2a) reporter only in β cells through Cre-mediated recombination under the control of the rat insulin promoter (RIP-Cre;Fucci2aR). Three-dimensional imaging of optically cleared pancreas tissue from these mice enabled the quantification of replicating β cells in islets and morphometric analysis of islets following mitogen treatment. Intravital imaging of RIP-Cre;Fucci2aR mice revealed cell cycle progression of β cells. Thus, this novel mouse line is a powerful tool for spatiotemporal analysis of β-cell proliferation in response to mitogen stimulation.

## Introduction

Diabetes is caused by β-cell dysfunction and deficiency. Stimulating β-cell proliferation is a promising treatment for diabetes, β as cells replicate very infrequently even in pathophysiological states. Many factors have been reported as potential β-cell mitogens, although the β-cell mitogenic effect by one of them has not been reproducible (Gusarova et al., 2014). The inconsistent results can be explained by the lack of a standardized method for quantifying replicating β cells (Cox et al., 2016). Thus far, determination of the β-cell proliferation rate has relied on immunohistochemical detection of cell cycle markers such as nucleotide analogs (5-bromo-2’-deoxyuridine [BrdU] and 5-ethynyl-2’-deoxyuridine [EdU]) or replication proteins (proliferating cell nuclear antigen and Ki-67). However, results obtained by the immunohistochemical assays show inter-laboratory variability (Cox et al., 2016) caused by methodological differences—e.g., in immunolabeling and image acquisition techniques. Replicating non-β cells within islets may also confound immunohistochemical analyses. Furthermore, there are presently no alternative methods that can be used to resolve these discrepant findings. Thus, new methods for quantifying replicating β cells are required in order to validate the effects of potential β-cell mitogens.

The fluorescent ubiquitination-based cell cycle indicator (Fucci) reporter is a well-known probe for monitoring cell cycle status (Sakaue-Sawano et al., 2008). The Fucci system relies on the expression of a pair of fluorescent proteins: mCherry-hCdt1(30/120) (a degron of chromatin licensing and DNA replication factor [Cdt] 1 fused to a fluorescent protein in the red spectrum) and mVenus-hGem (1/110) (a degron of Geminin fused to a fluorescent protein in the green spectrum). Reciprocal expression of these paired proteins labels cells in G_1_ phase and those in S/G_2_/M phase with red and green fluorescence, respectively. Thus, the Fucci system can be used to visualize the G_1_/S transition and quantify replicating cells.

In this study, we generated a mouse line in which only β cells expressing the Fucci probe are labeled according to cell cycle phase. Using this model, we specifically evaluated β-cell proliferation induced by administration of the insulin receptor antagonist S961 (a reported β-cell mitogen [Jiao et al., 2014]). In addition, we performed three-dimensional (3D) analyses of whole islets by observing optically cleared pancreas of these mice and found a strong and significant correlation between islet size and the number of replicating β cells per islet. These results demonstrate the utility of this mouse model for the study of β-cell proliferation.

## Results

### Generation of β cell-specific Fucci-expressing mice

To distinguish β cells in the G_0_/G_1_ phase from those in S/G_2_/M phase, we used Fucci technology, which is a proven tool for detecting actively proliferating cells. The R26-Fucci2aR transgenic mouse line harboring an upgraded Fucci2a reporter was recently generated in which Cre/loxP-mediated conditional expression of the Fucci2a transgene at the Rosa26 locus is driven by the cytomegalovirus early enhancer/chicken β actin promoter (Mort et al., 2014). By crossing rat insulin promoter (RIP)-Cre (Herrera et al., 1998) and R26-Fucci2aR mice, we generated the RIP-Cre;Fucci2aR line in which the Fucci2a probe is specifically expressed in β cells (Fig. 1A). RIP-Cre;Fucci2aR mice showed similar body weight and arbitrary blood glucose levels compared to Fucci2aR littermates (Fig. 1B and 1C), and there was no significant difference in blood glucose and insulin levels during in the oral glucose tolerance test (2 g/kg) between them (Fig. 1D and 1E), indicating that RIP-Cre; Fucci2aR mice have a normal metabolic profile.

**Figure 1.**
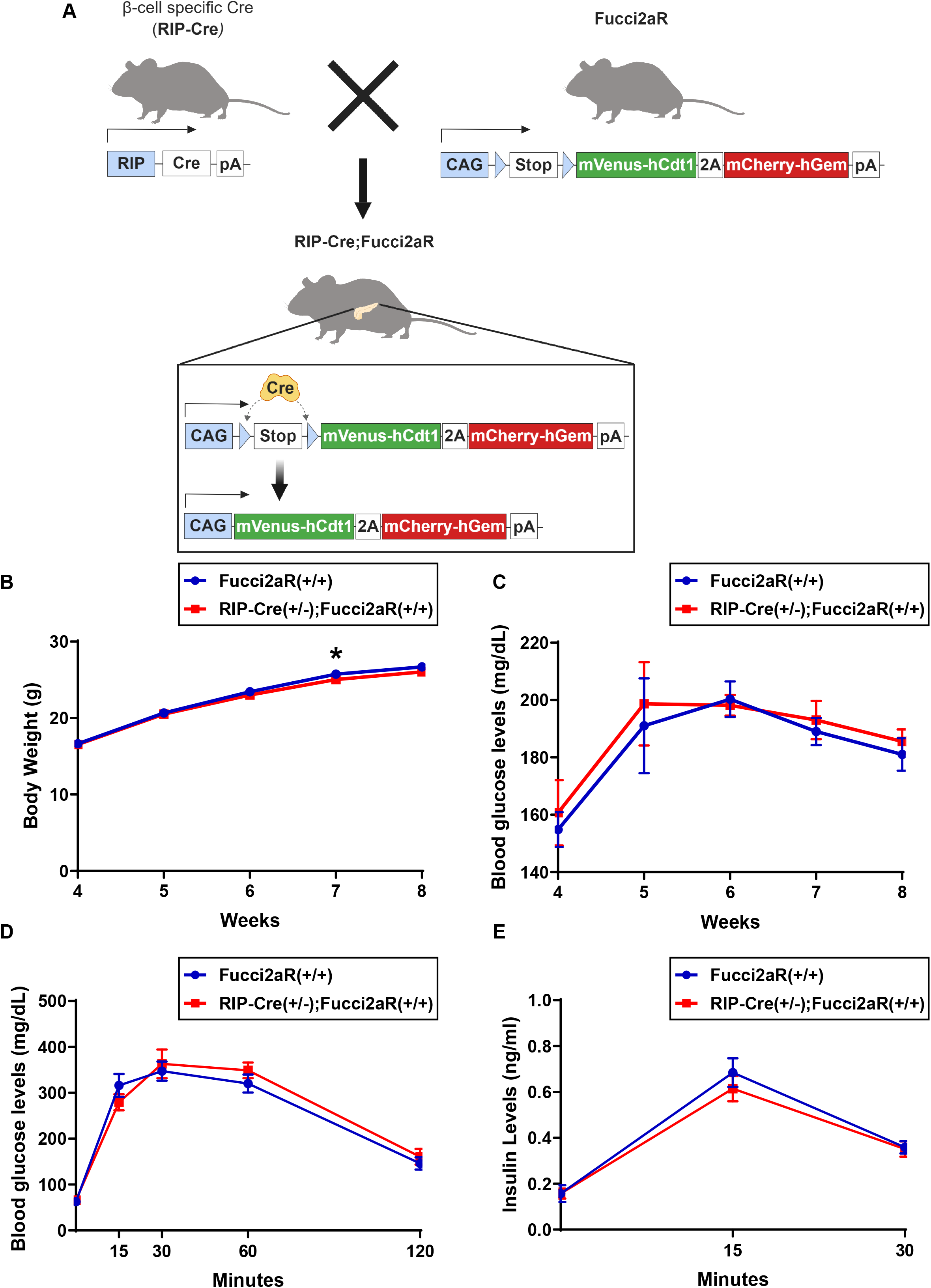
Genotype and metabolic phenotype of RIP-Cre;Fucci2aR mice. (A) Breeding scheme for the generation of RIP-Cre;Fucci2aR mice. RIP-Cre and Fucci2aR mouse lines were crossed to obtain RIP-Cre;Fucci2aR mice. After Cre-mediated recombination, the Fucci2a transgene was expressed specifically in β cells. (B) Body weight and (C) arbitrary blood glucose levels of RIP-Cre;Fucci2aR (N = 7) and Fucci2aR control (N = 7) mice during postnatal growth. (D, E) Oral glucose tolerance test (2 g/kg body weight) was performed on RIP-Cre; Fucci2aR (N = 7) and Fucci2aR control (N = 7) mice at 8 weeks. Data are expressed as mean ± SEM. *P < 0.05 (Mann-Whitney U test).

### β cell-specific Fucci expression in RIP-Cre;Fucci2aR mice

As proof of principle, we investigated the expression pattern of the Fucci2a probe in RIP-Cre;Fucci2aR mice. In order to characterize not only mCherry^+^ but also mVenus^+^ cells, we induced β-cell proliferation in RIP-Cre;Fucci2aR mice by continuous infusion of the vehicle phosphate-buffered saline (PBS) or insulin receptor antagonist S961 over 7 days with an osmotic pump. At the end of the treatment, frozen sections were prepared from the dissected pancreas and immunostained for insulin, glucagon, somatostatin, and Nkx-6.1, and the fluorescent signals of the Fucci2a probe were directly observed. In S961-treated RIP-Cre;Fucci2aR mice, mCherry and mVenus were expressed specifically in insulin^+^ and Nkx 6.1^+^ cells (Figure 2A and 2D), but not in glucagon^+^ or somatostatin^+^ cells (Figure 2B and 2C). To ensure that replicating β cells could be quantified, we compared the results of the EdU assay and the β-cell proliferation assay performed using RIP-Cre;Fucci2aR mice. Vehicle- and S961-treated mice were administered EdU 6 h before sacrifice, and mVenus^+^ cells (Figure 2E) or EdU^+^ insulin^+^ DAPI^+^ cells (Figure 2F) were counted in frozen sections. We found that the value of mVenus^+^ cells per β cells tended to be higher than that of EdU^+^ insulin^+^ cells per β cells.

**Figure 2.**
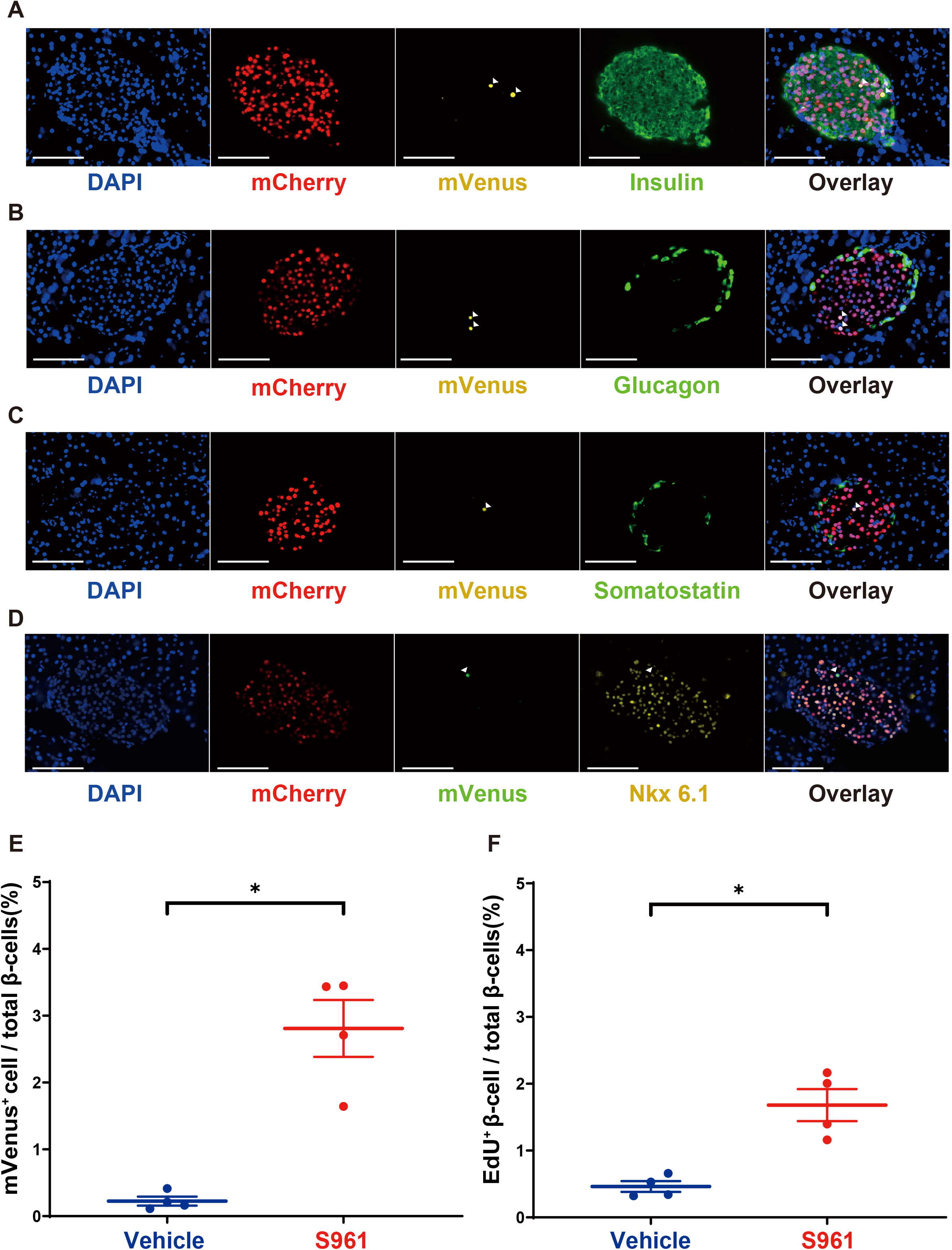
β Cell-specific expression of Fucci2a in RIP-Cre;Fucci2aR mice. (A–D) Frozen sections of pancreas tissue from RIP-Cre;Fucci2aR mice treated with S961 at 8 weeks of age immunostained for islet hormones and Nkx 6.1. Representative fluorescence images of mCherry^+^ (red) and mVenus^+^ (yellow) cells and immunofluorescence for islet hormones (green): insulin (A), glucagon (B), and somatostatin (C). (D) All mCherry^+^ (red) and mVenus^+^ (green) cells were Nkx 6.1-positive (yellow). Nuclei were stained with DAPI (blue). Scale bar, 100 μm. (E) Quantification of mVenus^+^ cells in the islets of RIP-Cre;Fucci2aR mice treated with vehicle (PBS; N = 4) or S961 (10 nM/week; N = 4). *P < 0.05. (F) Quantification of EdU+ β cells in islets of RIP-Cre;Fucci2aR mice treated with vehicle (PBS; N = 4) or S961 (10 nM/week; N = 4). *P < 0.05. Data are expressed as mean ± SEM.

### 3D Imaging of islets in RIP-Cre;Fucci2aR mice

Since each islet is densely packed with various cell types, replicating β cells can be misidentified in histological sections labeled for insulin and replication markers. In order to detect and quantify replicating β cells in 3D in whole islets of RIP-Cre; Fucci2aR mice, CUBIC clearing reagent (Kubota et al., 2017) was applied to pancreatic tissue samples from vehicle- or S961-treated RIP-Cre;Fucci2aR mice, and 3D images of the optically cleared tissue were obtained with a light sheet microscope equipped with a 5× objective lens. The spatial distributions of mVenus^+^ and mCherry^+^ cells were simultaneously visualized (Figure 3A–3F; Movie S1). Islets contained more mVenus^+^ cells following S961 treatment (Figure 3C and 3D). Spot objects corresponding to each mVenus^+^ or mCherry^+^ cells were reconstructed using Imaris Spot Detection and quantified by an automated process to determine the number of replicating β cells in each islet (Figure 3G and 3H). The diameter of β-cell cluster in each islet were measured using Imaris Surface tool. Thus, RIP-Cre;Fucci2aR mice are amenable to cross-sectional analyses of the number and spatial distribution of proliferating β cells.

**Figure 3.**
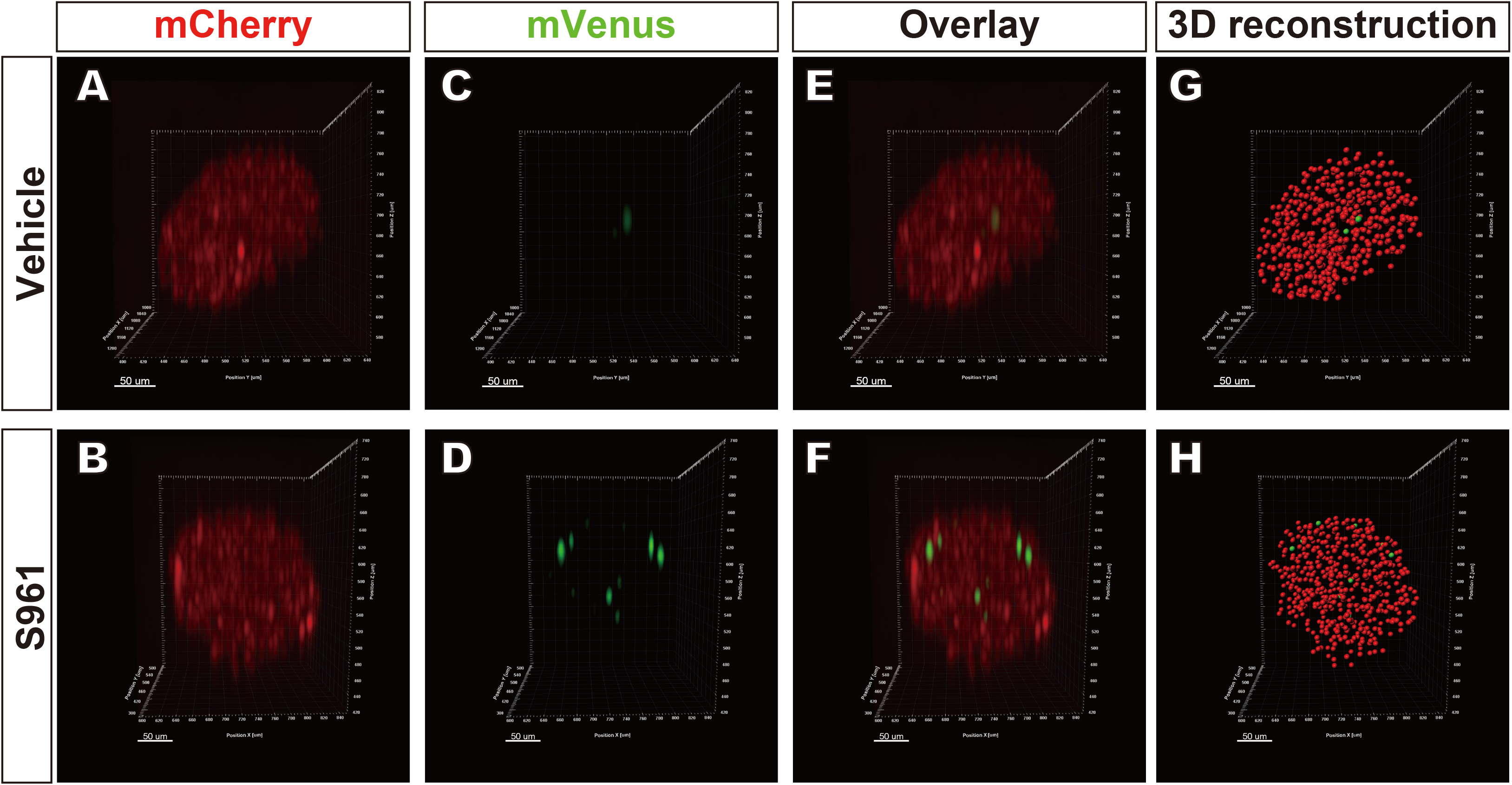
3D Imaging of islets in vehicle- and S961-treated RIP-Cre;Fucci2aR mice. Representative 3D images of islets following treatment for 1 week with vehicle or S961. (A, B) Representative fluorescence images of mCherry^+^ (red) and (C, D) mVenus^+^ (green) cells. (G, H) Morphological 3D reconstruction of mCherry^+^ (red) and mVenus^+^ (green) cells for automated cell counting. Images were obtained with a light-sheet microscope. Scale bar, 50 μm.

Given the utility of the Fucci2a probe for real-time monitoring of the cell cycle, we performed real-time in vivo imaging in S961-treated RIP-Cre;Fucci2aR mice using a two-photon microscope equipped with a 25× water objective lens. This intravital imaging of an islet in a RIP-Cre;Fucci2aR mouse initiated 40 h after S961 treatment revealed the G_1_-S transitions of β cells (Supplemental Information Movies S2).

### The number of replicating β cells per islet is positively correlated with islet size

The relationship between the number of replicating β cells per islet and the morphological characteristics of islets is unclear. We address this issue by analyzing 3D images obtained from RIP-Cre;Fucci2aR mice. Blood glucose and insulin levels were higher in mice treated with S961 (N = 4) than in those treated with vehicle (N = 4) (Figure 4A, 4B). When we examined all islets whose β-cell cluster diameter was over 100 μm, the β-cell cluster diameter and β-cell number per islet were greater in S961-treated RIP-Cre; Fucci2aR mice (Figure 4C, 4D, 4E). In addition, the proportion of mVenus^+^ cells per islet were higher in S961-treated as compared to control mice (Figure 4F). Moreover, the mVenus^+^ cell number per islet was positively correlated with β-cell number per islet in both vehicle-treated (Figure 4G; r = 0.87, P < 0.0001) and S961-treated (Figure 4G; r = 0.84, P < 0.0001) mice.

**Figure 4.**
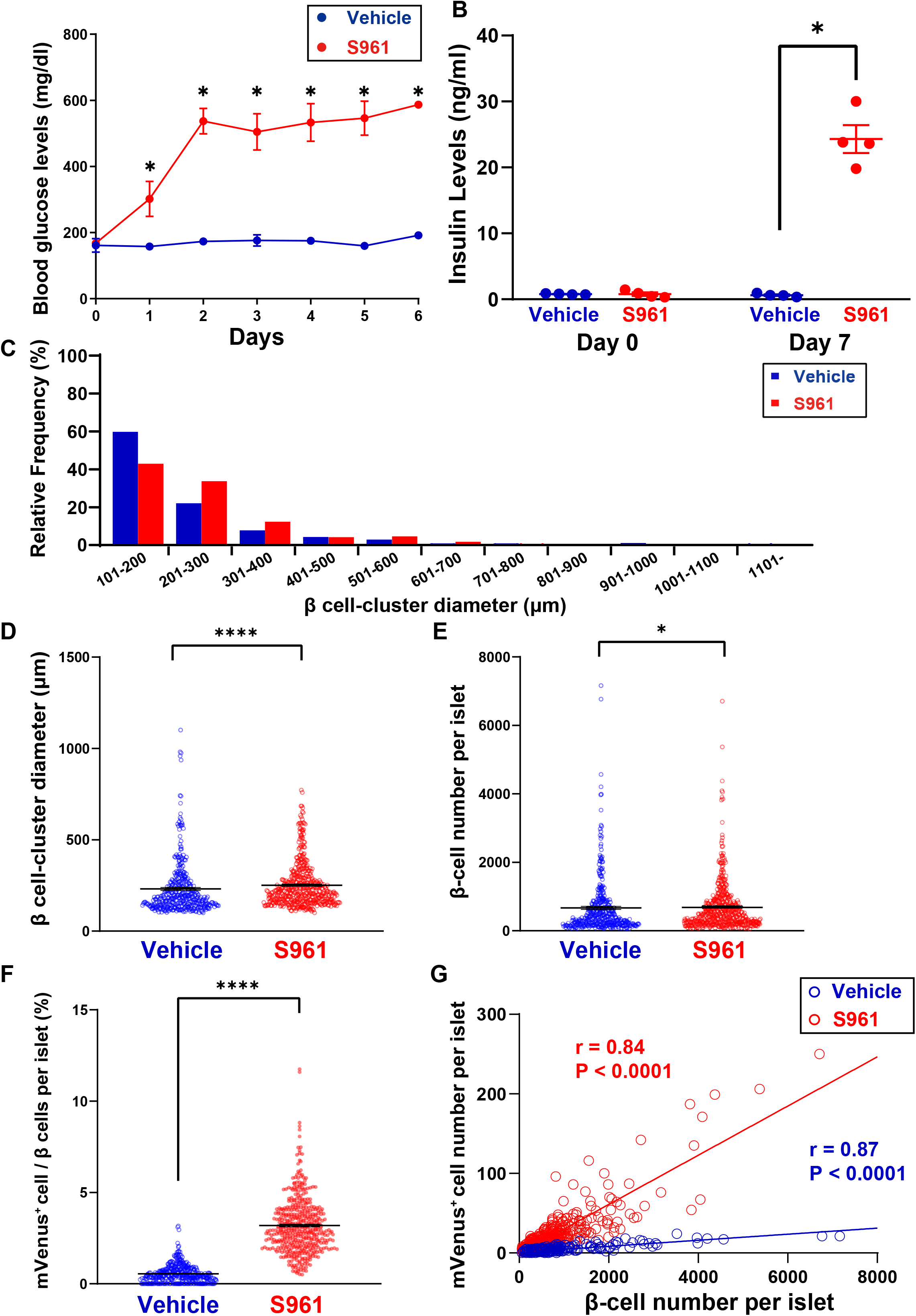
Quantification of replicating β cells in RIP-Cre;Fucci2aR mice following S961 treatment. (A, B) RIP-Cre;Fucci2aR mice were treated with S961 (10 nM/week; N = 4) or vehicle (PBS; N = 4) for 7 days. (A) Arbitrary blood glucose and (B) serum insulin levels at the end of the 7-day treatment. (C) Histogram of β-cell cluster diameter in S961- and vehicle-treated RIP-Cre;Fucci2aR mice. Morphometric analysis was performed on islets harboring β-cell clusters with a diameter > 100 μm (S961, n = 454 islets, N = 4 mice; vehicle, n = 348 islets, N = 4 mice). (D) β-cell cluster diameter in S961- and vehicle-treated mice. ****P < 0.0001 (N = 4). (E) Number of β cells per islet in S961- or vehicle-treated mice. *P < 0.05 (N = 4). (F) Percentage of mVenus^+^ cells per islet in S961- and vehicle-treated mice. ****P < 0.0001 (N = 4). (G) Correlation between number of mVenus^+^ cells and number of β cells per islet. mVenus^+^ cell number and β cell number per islet was strongly correlated in both groups (S961, r = 0.87, P < 0.0001; vehicle, r = 0.77, P < 0.0001). Data are presented as mean ± SEM.

Next, we investigated whether this positive correlation could be also found under a physiological condition such as diet-induced obesity. The RIP-Cre;Fucci2aR mice were divided into two groups: one fed with high-fat diets (HFD) and the other fed with control diets for 13 weeks. HFD group (N = 7) gained significantly more body weight than control diet group (N = 7) at the end of 13-week feedings (Figure 5A). Compared to control diet group, HFD group revealed greater β-cell cluster diameter (Figure 5B, 5C), more β-cell number per islet (Figure 5D) and higher proportion of mVenus^+^ cells per islet (Figure 5E). Finally, the positive correlation between mVenus^+^ cell number per islet and β-cell number per islet was also found in both HFD group (Figure 5F; r = 0.81, P < 0.0001) and control diet group (Figure 5F; r = 0.60, P < 0.0001). These data indicate that the number of replicating β cells per islet depends on the size of the islet.

**Figure 5.**
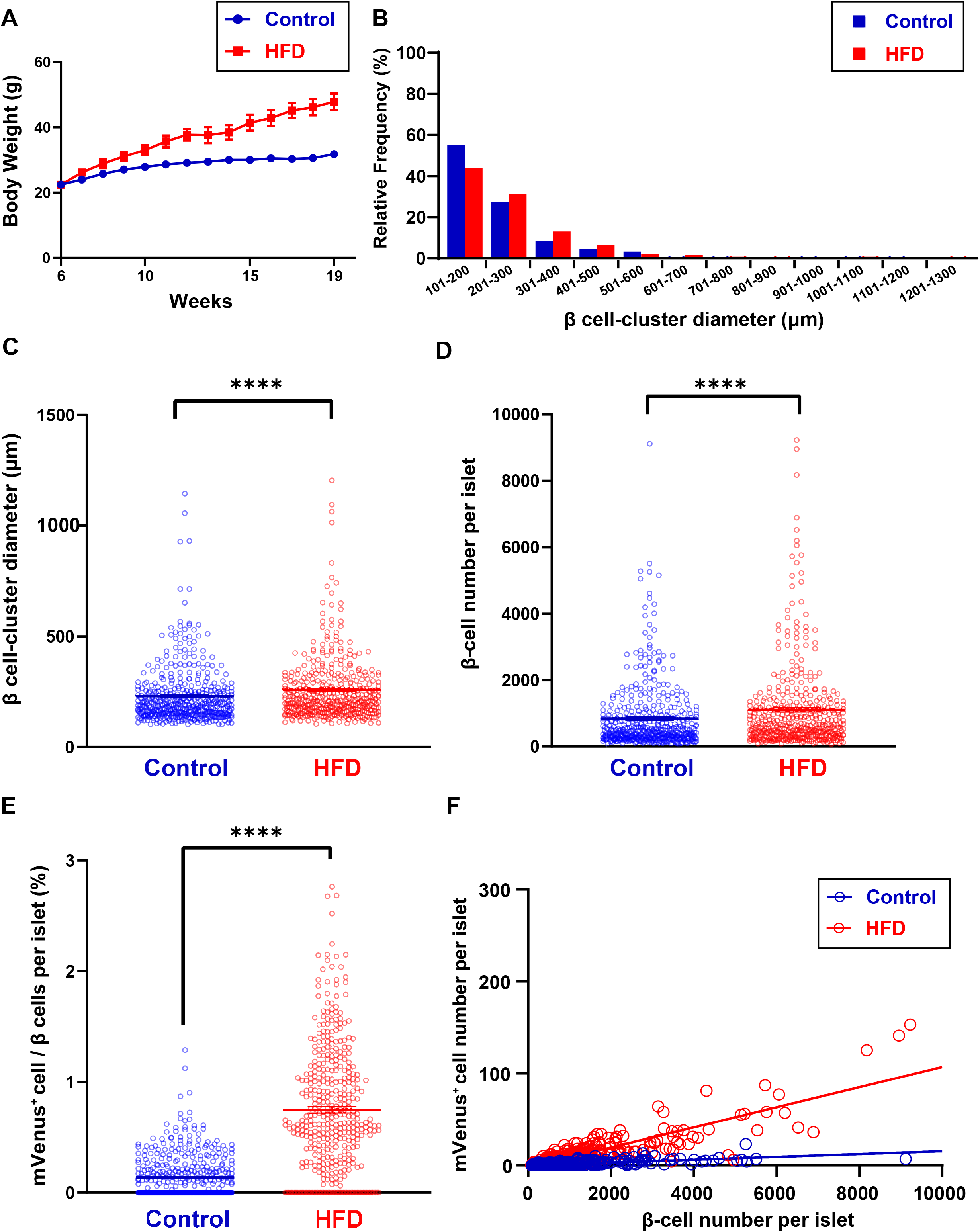
Quantification of replicating β cells in RIP-Cre;Fucci2aR mice following high-fat diet feeding. (A) Body weight of RIP-Cre;Fucci2aR mice fed with high-fat diets (HFD; N = 7) or control diets (Control; N = 7) for 13 weeks. (B) Histogram of β-cell cluster diameter in RIP-Cre;Fucci2aR mice under HFD or control diet feeding. Morphometric analysis was performed on islets harboring β-cell clusters with a diameter > 100 μm (HFD, n = 407 islets, N = 4 mice; Control, n = 432 islets, N = 4 mice). (C) β-cell cluster diameter. ****P < 0.0001 (N = 4). (D) Number of β cells per islet. ****P < 0.0001 (N = 4). (E) Percentage of mVenus^+^ cells per islet. ****P < 0.0001 (N = 4). (F) Correlation between number of mVenus^+^ cells and number of β cells per islet. mVenus^+^ cell number and β cell number per islet was strongly correlated in both groups (HFD, r = 0.81, P < 0.0001; Control, r = 0.60, P < 0.0001). Data are presented as mean ± SEM.

### β-cell proliferation induced by S961 is not due to hyperglycemia

Hyperglycemia has been shown to induce β-cell proliferation (Alonso et al., 2007; Porat et al., 2011). Since S961 administration causes hyperglycemia, we investigated whether this mediates β-cell proliferation during S961 treatment. To exclude the effects of hyperglycemia, we normalized blood glucose levels in S961-treated RIP-Cre;Fucci2aR mice by co-administration of sodium–glucose cotransporter 2 inhibitor (SGLT2i). The mice were divided into four groups: vehicle treatment with control diet (vehicle + control), vehicle treatment with control diet containing 0.02% empagliflozin (vehicle + 0.02% empagliflozin), S961 treatment with control diet (S961 + control), and S961 treatment with control diet containing 0.02% empagliflozin (S961 + 0.02% empagliflozin). Although blood glucose level was higher in the S961 + control than in the S961 + 0.02% empagliflozin group, the level in the latter was similar to that in the vehicle + control group (Figure 6A). The S961 + 0.02% empagliflozin group had a lower insulin level than the S961 + control group but nonetheless showed hyperinsulinemia (Figure 6B), reflecting the continuous pharmacological action of S961 irrespective of empagliflozin co-administration.

**Figure 6.**
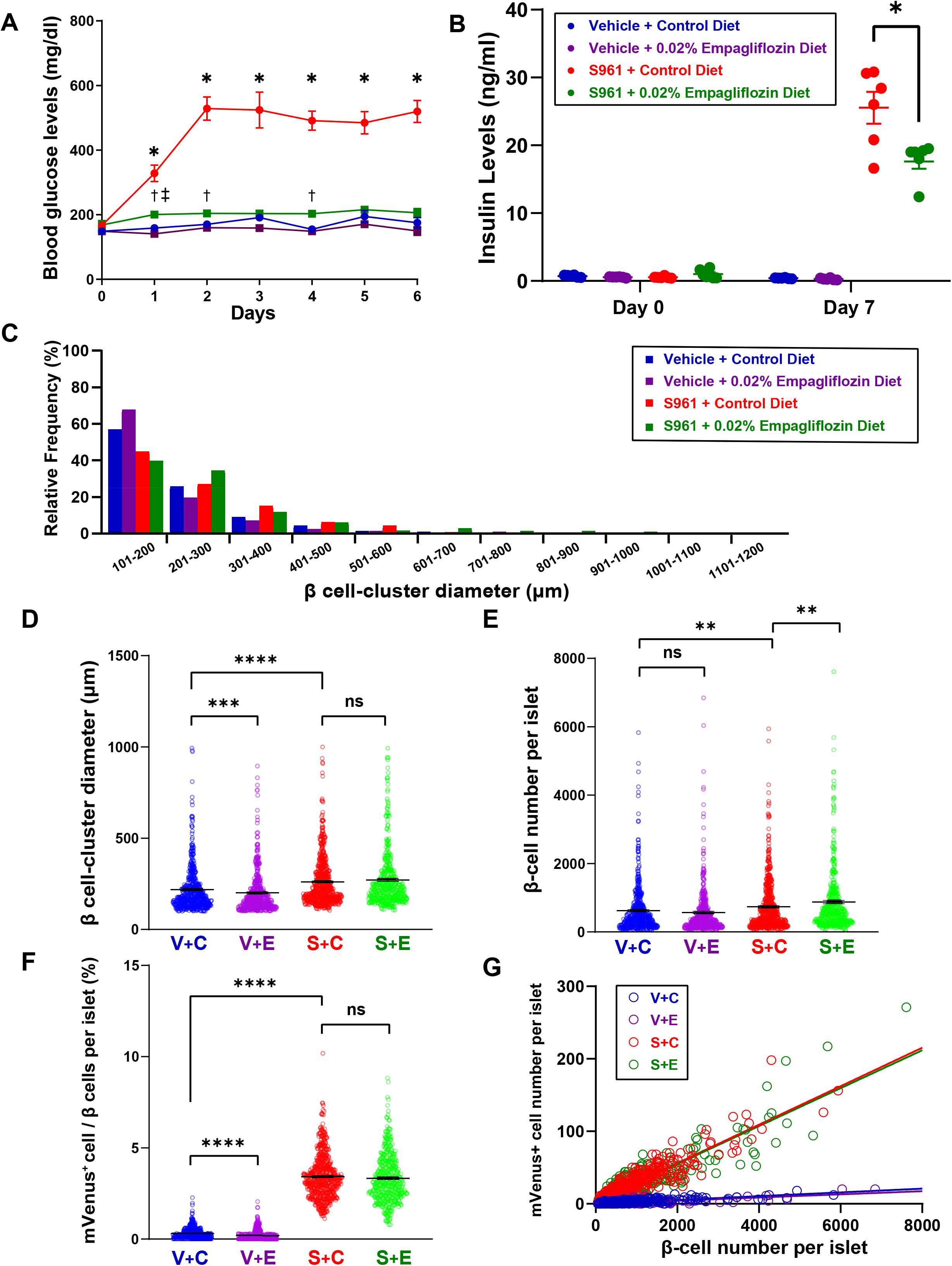
S961-induced β cell proliferation is not mediated by hyperglycemia. (A, B) RIP-Cre;Fucci2aR mice were divided into four groups: 1) vehicle + control, treated with vehicle and fed a control diet (N = 6); 2) vehicle + 0.02% empagliflozin, treated with vehicle and fed a diet supplemented with 0.02% empagliflozin (N = 6); 3) S961 + control, treated with S961 (10 nM/week) and fed a control diet (N = 6); and 4) S961 + 0.02% empagliflozin, treated with S961 (10 nM/week) and fed a diet supplemented with 0.02% empagliflozin (N = 6). (A) Arbitrary blood glucose levels. *P < 0.05, S961 + control diet group vs. S961 + 0.02% empagliflozin group. ^†^P < 0.05, S961 + 0.02% empagliflozin group vs. Vehicle + control group. ^‡^P < 0.05, vehicle + control group vs. vehicle + 0.02% empagliflozin group. (B) Serum insulin levels at the end of the 7-day treatment. *P < 0.05, S961 + control group vs. S961 + 0.02% empagliflozin group. (C) Histogram of β-cell cluster diameter. Morphometric analysis was performed on islets harboring β-cell clusters with a diameter > 100 μm (vehicle + control, N = 4 mice, n = 496 islets; vehicle + 0.02% empagliflozin, N = 4 mice, n = 440 islets; S961 + control, N = 4 mice, n = 544 islets; S961 + 0.02% empagliflozin, N = 4 mice, n = 391 islets). (D) β-cell cluster diameter, (E) number of β cells per islet, and (F) percentage of mVenus^+^ cells per islet. **P < 0.01 (N = 4), ***P < 0.001 (N = 4), ****P < 0.0001 (N = 4); ns, not significant. (G) Correlation between number of mVenus^+^ cells and number of β cells per islet. mVenus^+^ cell and β-cell number per islet were strongly correlated in all groups (vehicle + control, r = 0.70, P < 0.0001; vehicle + 0.02% empagliflozin diet, r = 0.61, P < 0.0001; S961 + control diet, r = 0.93, P < 0.0001; and S961 + 0.02% empagliflozin diet, r = 0.88, P < 0.0001). V + C, vehicle + control; V + E, vehicle + 0.02% empagliflozin diet; S + C, S961 + control diet; S + E, S961 + 0.02% empagliflozin diet. Data are presented as mean ± SEM.

We next investigated the morphological characteristics of islets in all four groups and found that islets were larger in mice treated with S961 as compared to vehicle (Figure 6C, 6D, 6E). β-cell cluster diameter was larger in the vehicle + control as compared to the vehicle + 0.02% empagliflozin group, while no difference was observed between S961 + control and S961 + 0.02% empagliflozin groups (Figure 6C, 6D). On the other hand, while the number of β cells per islet was greater in S961 + 0.02% empagliflozin group as compared to S961 + control group (Figure 6E), there was no difference in the proportion of mVenus^+^ cells number per islet between the two groups (Figure 6F). In all four groups, there were positive correlations between mVenus^+^ cell number and β-cell number per islet (vehicle + control: r = 0.78, P < 0.0001; vehicle + 0.02% empagliflozin diet: r = 0.74, P < 0.0001; S961 + control diet: r = 0.92, P < 0.0001; and S961 + 0.02% empagliflozin diet: r = 0.90, P < 0.0001; Figure 6G). These results indicate that hyperglycemia does not mediate S961-induced β-cell replication.

## Discussion

Identifying potential β-cell mitogens could lead to a novel diabetes therapy. Although many factors that control β-cell replication have been identified to date, their mitogenic effects on β cells have not been precisely evaluated since immunohistochemical assays are unreliable for accurately identifying β cells. A recent study reported inter-laboratory variability in the immunohistochemical detection of Ki-67 for the identification of β cells and quantification of their replication; the authors concluded that the discrepancy among laboratories was due to the misidentification of replicating non-β cells within islets as β cells (Cox et al., 2016). Since many different cell types are densely packed in sphere-like islets, the analysis of 2D immunohistochemical data could account for inaccuracies in the detection of β cells. Furthermore, the nucleotide analog BrdU, which is often used as a cell cycle marker in traditional immunohistochemical assays, has unfavorable effects on the cell cycle of β cells (de Casteele et al., 2013). Thus, immunohistochemistry using cell cycle markers such as Ki-67 and BrdU are not appropriate for evaluating β-cell proliferation. In order to overcome these limitations, we developed a novel assay that can precisely detect and be used to quantify replicating β cells.

Fucci2a, a single fluorescence marker for monitoring cell cycle transition, differentiates cells in G_0_/G_1_ from those in S/G_2_/M phase based on mCherry-hCdt1 (30/120) and mVenus-hGem (1/110) expression (Sakaue-Sawano et al., 2008). Using RIP-Cre;Fucci2aR mice in which Fucci2a is expressed specifically in β cells, we established an assay for detecting the proliferative β-cell pool that is uncontaminated by replicating non-βcells. Our results also suggest that the Fucci2a assay is more sensitive than the short-term EdU labeling assay, which cannot detect β cells in G_2_/M phase.

3D Analysis of RIP-Cre/Fucci2aR mice allows more precise evaluation of replicating β cells, and increases the sample size compared to 2D immunohistochemical assays. We optically cleared pancreas tissue samples from RIP-Cre/Fucci2aR mice for 3D fluorescence imaging, which provided spatial information on replicating β cells within each islet. This 3D analysis allowed us to examine the correlation between β-cell proliferative capacity and the morphological characteristics of each islet. Furthermore, intravital imaging demonstrated that longitudinal spatiotemporal data on β-cell proliferation can be obtained from RIP-Cre/Fucci2aR mice. The 3D analysis of the pancreas of RIP-Cre/Fucci2aR mice revealed a higher β-cell proliferation rate within each islet in mice treated with S961 than in those treated with vehicle. When we counted all mCherry^+^ and mVenus^+^ cells as β cells, we found that the total number of β cells per islet was increased by S961 treatment. This is consistent with previous reports on S961-induced β cell proliferation and mass expansion (Jiao et al., 2014). In addition, the strong positive correlation between mVenus^+^ cell and total β-cell number per islet suggested that larger islets contain more replicating β cells.

The signals that regulate β-cell proliferation upon S961 treatment are not known. Hyperglycemia has been reported to directly induce β-cell proliferation, while a recent study showed that S961-induced hyperinsulinemia and β cell mass expansion occurred even when blood glucose levels were normalized by co-administration of a monoclonal glucagon receptor antibody (Okamoto et al., 2017). Our results showed that S961 stimulated β-cell proliferation and increased β-cell number per islet even when hyperglycemia was normalized by SGLT2i treatment. These results provide new evidence for the existence of mitogenic factors mediating S961-induced β-cell proliferation, except under hyperglycemic conditions.

Our study had several limitations. Firstly, since only β cells were labeled by the Fucci2a probe, other endocrine cells within islets could not be detected in RIP-Cre;Fucci2aR mice. Therefore, mitogenic effects on non-β cells must be investigated using other methods. Secondly, the attenuation of fluorescence by light scattering limited the observation depth from the pancreas surface. Such signal attenuation is unavoidable despite the optical clearing process. Under the light sheet microscope, only islets near (~2.0 mm from) the surface were clearly detected for quantification of fluorescent cells. Although this restricts the size of the islet population, the sample size is still larger using our method as compared to a conventional immunohistochemical assay because it is based on 3D analysis of the whole pancreas.

In summary, the transgenic mouse line expressing the Fucci2a probe in β cells developed in this study provides a new tool for analyzing β-cell proliferation in a more reliable and reproducible manner than conventional immunohistochemistry. The high spatial resolution of the 3D images obtained with a light-sheet microscope allows accurate detection of replicating β-cells. This system can be useful for validating the efficacy and therapeutic potential of β-cell mitogens for inhibition of diabetes progression.

## Acknowledgments

The authors thank Asako Sakaue-Sawano and Atsushi Miyawaki for their scientific discussion; Saki Kanda and Sara Yasui for technical assistance; and Yukiko Inokuchi, Yukiko Tanaka, and Fumiko Uwamori for secretarial assistance. This work was supported by Kyoto University Live Imaging Center and in part by Grants-in-Aid KAKENHI 16H06280 “ABiS”.

## Author Contributions

The study was designed by S.T., D.Y., and N.I. Experiments were performed by S.T. and A.B., and data were analyzed by S.T. The manuscript was written by S.T., D.Y., and N.I. with input from all authors.

## STAR Methods

### CONTACT FOR REAGENT AND RESOURCE SHARING

Further information and requests for resources and reagents should be directed to and will be fulfilled by the Lead Contact, Nobuya Inagaki. RIP-Cre mice were obtained under Material Transfer Agreements from Prof. Pedro L. Herrera. Fucci2aR mouse strain (RBRC06511) was provided by RIKEN BRC through the National BioResource Project of the MEXT/AMED, Japan.

### EXPERIMENTAL MODEL AND SUBJECT DETAILS

#### Generation of the transgenic mouse line

To establish the mouse model for studying β-cell proliferation, we used mice harboring R26-Fucci2aR (RIKEN BRC through Kyoto University Medical Science and Business Liaison Organization). This newer Fucci2a reporter is a bicistronic Cre-inducible probe consisting of two fluorescent proteins: truncated Cdt1 (30/120) fused to mCherry, and truncated Geminin (1/110) fused to mVenus. The two fusion proteins are always alternately expressed according to cell cycle phase in the same ratio, making it possible to detect and quantify the number of labeled cells. By crossing Rip-Cre and Fucci2aR mice, we generated RIP-Cre;Fucci2aR mice expressing the Fucci2a reporter in a β cell-specific manner. In these RIP-Cre;Fucci2aR mice, mCherry-hCdt1 (30/120) (red fluorescence) and mVenus-hGem (1/110) (green fluorescence) are expressed in β cell nuclei during G0/G1 and S/G2/M phases, respectively. Only hemizygous males on the C57BL/6 background were used in this study. Mice had free access to standard rodent chow and water and were housed in a temperature-controlled environment under a 14:10-h light/dark cycle. Animal care and protocols were reviewed and approved by the Animal Care and Use Committee of Kyoto University Graduate School of Medicine (MedKyo15298).

### METHOD DETAILS

#### In vivo mouse studies

S961 was obtained from Novo Nordisk (Bagsværd, Denmark). Vehicle (PBS) or 10 nmol S961 was loaded into an osmotic pump (Alzet 2001; DURECT Corp., Cupertino, CA, USA) subcutaneously implanted into the back of RIP-Cre;Fucci2aR mice at 8 weeks of age. Mice were euthanized and the pancreas was harvested 7 days after S961 or vehicle treatment.

Blood glucose levels were measured daily. Plasma was collected on days 0 and 7 to measure insulin level. For a model of diet-induced obesity, six-week old RIP-Cre;Fucci2aR mice were fed with high-fat diets (Research Diet, cat. no. D12492) or control diets (Research Diet, cat. no. D12450J) for 13 weeks, and body weight were measured weekly. For the EdU labeling assay, mice were intraperitoneally injected with EdU (50 mg/kg) 6 h before sacrifice.

#### Oral glucose tolerance test

Mice were fasted for 16 h and then orally administered a 20% glucose solution (2 g/kg body weight). Blood samples were collected from the tail vein of mice 0, 15, and 30 min after glucose loading using heparinized calibrated glass capillary tubes (cat. no. 2-000-044-H; Drummond Scientific Co., Broomall, PA, USA). Blood glucose level was measured using the Glutest Neo Sensor (Sanwa Kagaku Kenkyusho, Nagoya, Japan). Plasma samples were prepared by centrifuging the blood samples at 9000 × *g* for 10 min, and insulin level was measured using the Ultra Sensitive PLUS Mouse Insulin ELISA kit (cat. no. 49170-53; Morinaga, Tokyo, Japan).

#### Immunohistochemical observation of tissue sections

Mice were anesthetized by intraperitoneal injection of pentobarbital sodium (10 mg/kg); a 26-gauge needle was inserted into the left ventricle through the apex, and the mice were transcardially perfused with cold PBS followed by cold 4% paraformaldehyde (PFA, Wako Pure Chemical Industries, Osaka, Japan). The harvested pancreas was immediately immersed in PFA at 4°C with gentle shaking for less than 24 h, and then embedded in Optimal Cutting Temperature compound. Frozen samples were cut into 8-μm sections. After air drying, the frozen sections were incubated with blocking buffer composed of PBS with 10% (v/v) goat serum and 0.2% (v/v) Triton-X100) for 30 min at room temperature, and then incubated overnight at room temperature in blocking buffer supplemented with rabbit anti-insulin (200-fold dilution; cat. no. ab181547), mouse anti-glucagon (2000-fold dilution; cat. no. ab10988), rat anti-somatostatin (100-fold dilution; cat. no. ab30788), or rabbit anti-Nkx 6.1 (100-fold dilution; cat. no. ab221549) antibody (all from Abcam, Cambridge, MA, USA), followed by Alexa Fluor 647-conjugated goat anti-rabbit IgG (H+L) (200-fold dilution; cat. no. A-21245; Thermo Fisher Scientific, Waltham, MA, USA), Alexa Fluor 647-conjugated goat anti-mouse IgG (H+L) (200-fold dilution; cat. no. ab150115; Abcam), or Alexa Fluor 647-conjugated goat anti-rat IgG (H+L) (200-fold dilution; cat. no. ab150159; Abcam) for 1 h at room temperature. The sections were incubated in PBS containing DAPI (final concentration: 0.01 mg/ml) for 15 min at room temperature and mounted with Vectashield (Vector Laboratories, Burlingame, CA, USA) on 24 × 40-mm coverslips (cat. no. C024401; Matsunami Glass, Osaka, Japan). Immunolabeled tissue sections were observed with an inverted fluorescence microscope (BZ-X710; Keyence, Osaka, Japan). EdU was detected using the Click-iT EdU Alexa Fluor 647 kit (Thermo Fisher Scientific) according to the manufacturer’s protocol.

#### Tissue clearing

Pancreas tissue samples were collected and fixed as described above, washed three times for more than 2 h each time in PBS at room temperature with gentle shaking. For delipidation and permeabilization, the samples were immersed in 50% (v/v) CUBIC-L clearing reagent for at least 6 h followed by CUBIC-L at 37°C with gentle shaking for 3 days. The CUBIC-L reagent was refreshed daily during this period. After clearing, samples were immersed in 50% (v/v) CUBIC-R for at least 6 h and in CUBIC-R at room temperature with gentle shaking for at least 2 days.

#### 3D Imaging

3D Images of optically cleared pancreas tissue were acquired with a light-sheet microscope (Lightsheet Z.1; Carl Zeiss, Oberkochen, Germany) equipped with a 5×/0.16 NA objective lens. For mCherry-hCdt1 (30/120) imaging, we used 22% laser power (561-nm laser) and a 28-ms exposure time. For mVenus-hGem (1/110) imaging, we used 90% laser power (488-nm laser) and a 70-ms exposure time. The z-stack images (1920 × 1920 pixel, 16-bit) were acquired at 4.63 μm intervals.

#### Intravital imaging

Following S961 treatment for 40 h, RIP-Cre;Fucci2aR mice were anesthetized by 1.5%–2% isoflurane (Wako Pure Chemical Industries) inhalation. The hair on the abdominal area was removed and skin disinfected with 70% ethanol. A small oblique incision running parallel to the last left rib was made to expose the pancreas on the left side of the abdominal wall. The mice then were placed in the supine position on an electric heating pad maintained at 37°C. The pancreas was immobilized using a suction imaging device (Figure S1), and time-lapse imaging was performed with a two-photon excitation microscope (FV1200MPE-BX61WI; Olympus, Tokyo, Japan) equipped with a 25×/1.05 NA water-immersion objective lens (XLPLN 25XWMP; Olympus) and an In-Sight DeepSee Ultrafast laser (Spectra Physics, Santa Clara, CA, USA). Images were acquired every 5 min for ~10 h in 5-μm steps at a scan speed of 20 μs/pixel. Mice were euthanized after imaging.

#### Image processing

Acquired images were analyzed with the 3D reconstruction software Imaris (Bitplane AG, Zurich, Switzerland). A whole series of consecutive 2D cross-sectional images was reconstructed into a 3D structure using the “Volume rendering” function. Each islet was then isolated using the “Crop 3D function”, and a Gaussian filter was applied for background noise reduction. A spot detection algorithm was used for automated cell identification and cell counts. Morphometric measurements of maximum diameter of β cells within each islet were obtained using the Imaris surface creation tool.

#### Quantification and statistical analysis

The Mann–Whitney U test was performed to evaluate the difference between two sets of data. P values < 0.05 were considered statistically significant. No statistical method was used to predetermine sample size. Statistical analyses were performed using GraphPad Prism (GraphPad Software, La Jolla, CA, USA).

## Supplemental Information

**Figure S1.** Experimental setup of intravital pancreas imaging by two-photon microscopy, Related to STAR Methods section.

**Movie S1.** 3D Imaging of Islets in RIP-Cre;Fucci2aR mice, Related to Figure 3 The movie shows how 3D images of islets were edited and analyzed using Imaris software.

**Movie S2.** *In vivo* imaging of an islet in a RIP-Cre;Fucci2aR mouse, Related to Figure 3 The movie shows the G1-S transitions in two β cells, which are also automatically detected using the cell-tracking tool in Imaris. Scale bars, 50 μm. Time is shown in hours:minutes.

