## Supplemental Figure 1 for "Three-dimensional analysis of β-cell proliferation by a novel mouse model"

**Figure S1**

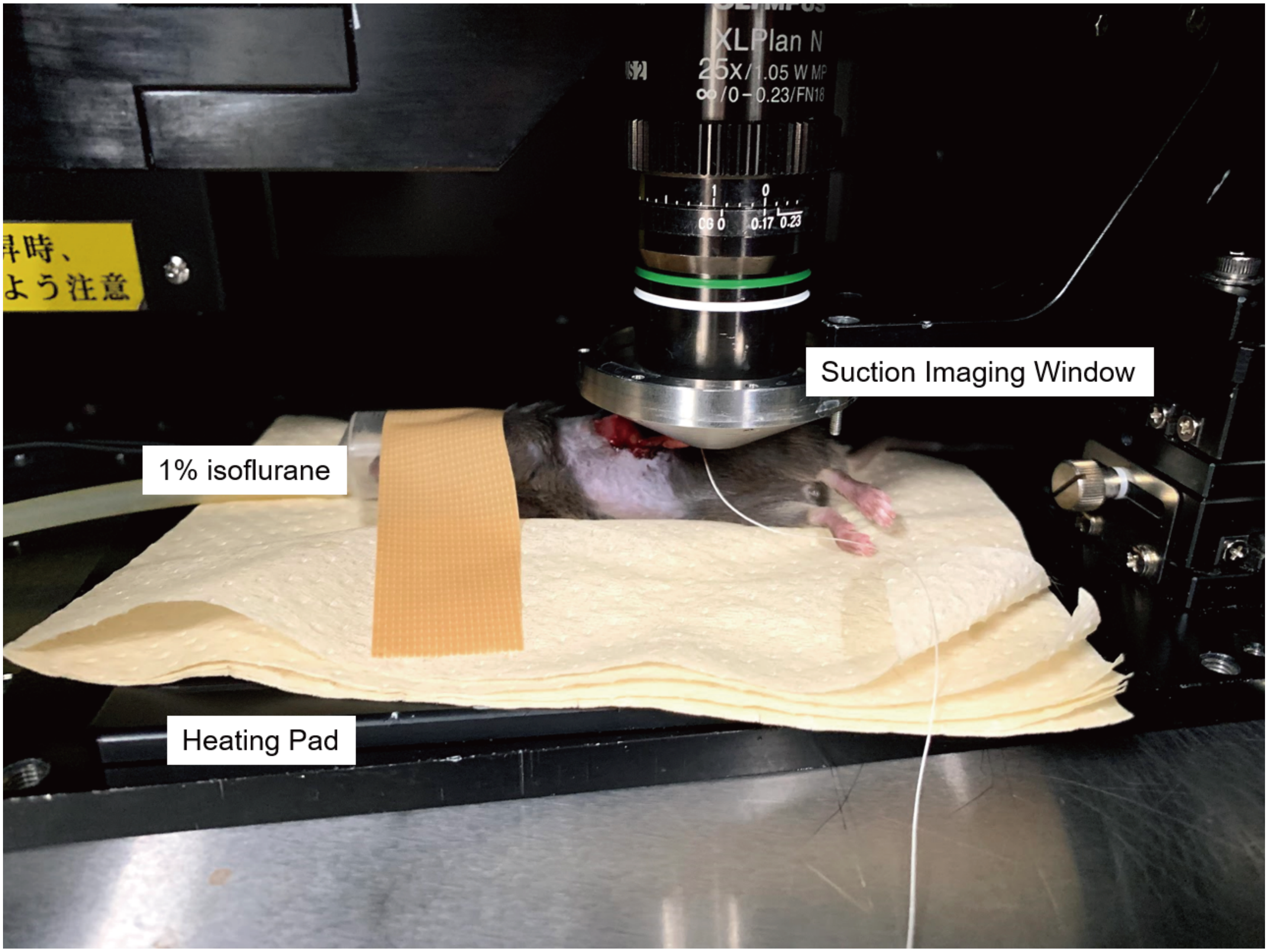


**Figure S1.** Experimental setup of intravital pancreas imaging by two-photon microscopy, Related to STAR Methods section.
